# Antifungal tolerance and resistance dynamics depend on infection load, drug treatment, and host innate immunity

**DOI:** 10.64898/2026.06.04.730223

**Authors:** Clare Maristela V. Galon, Daniel A. Charlebois

**Affiliations:** Department of Physics, University of Alberta, 11455 Saskatchewan Drive NW, Edmonton, Alberta, T6G 2E1, Canada

**Keywords:** Antifungal tolerance, antimicrobial resistance evolution, *Candidozyma auris*, *Galleria mellonella*, innate host immunity, population dynamics models

## Abstract

Fungi contribute substantially to the global antimicrobial resistance crisis. Tolerance is a novel form of antifungal resistance in which fungal pathogens emerge, grow slowly, and survive antifungal drug treatment. We quantitatively model the population and evolutionary dynamics of a multidrug resistant pathogen *Candidozyma auris* (formerly *Candida auris*) infecting an invertebrate host (*Galleria mellonella*) with an innate immune system. We find that the establishment, dominance, and co-existence of tolerant and resistant subpopulations depend on infection load, drug-treatment, and innate host immunity. This study enhances our understanding of the population and evolutionary dynamics of pathogenic infections and provides a quantitative framework to predict antimicrobial resistance.

## 1. Introduction

Increasing mortality rates due to antimicrobial resistance (AMR) is a growing threat to healthcare systems worldwide [1, 2]. Infectious fungal diseases (IFDs) are on the rise and are attributed to approximately 150 million cases and 1.7 million deaths across the globe each year [3]. Climate change is expected to further exacerbate this problem, as potentially pathogenic fungi adapt to rising temperatures, expand their geographic range, and bypass the thermal defenses of warm-blooded hosts [4]. Among the pathogenic fungi contributing to infectious fungal diseases (IFDs) and antifungal resistance (AFR), is the multidrug resistant human fungal pathogen *Candidozyma auris* (*C. auris*). This pathogen was classified in the critical priority group of the World Health Organization Fungal Pathogens Priority List [1], *C. auris* is associated with mortality rates ranging from 40 % to 70 % for bloodstream infections [5, 6], and in rare cases, pandrug resistance [7, 8]. This emerging healthcare crisis is compounded by the near-absence of new antifungal drugs [9, 10] and a lack of rapid, non-culture based diagnostics [11].

Recently, antifungal tolerance has been identified as an important factor contributing to AFR. Tolerance occurs when fungi emerge slowly during drug treatment above the minimum inhibitory concentration (MIC) of an antifungal drug [12, 13, 14]. Tolerance is a reversible phenomenon, in that pathogenic yeast can switch between drug-susceptible and drug-tolerant phenotypes [15]. It has also been established that drug concentration differentially selects for tolerance and resistance [16]. While the mechanisms underlying tolerance are still being elucidated, tolerance has been associated with stress responses and is thought to be affected by many pathways [17, 18, 19]. The failure of antifungal treatment may not always be driven by genetic resistance, but by transient tolerance that allows fungal pathogens to survive long enough to evolve permanent resistance within the host.

Quantitative modeling studies have predicted that non-genetic resistance enhances the survival of invading microbial populations during drug treatment [20, 21], while at the same time slowing the fixation of permanent genetic resistance [22]. This occurs because susceptible, non-genetically resistant, and genetically resistant subpopulations compete for shared resources, impeding the spread of genetic mutants through the infecting population. These dynamics were observed for both static antifungal drugs, which slow fungal growth, and cidal drugs, which kill fungi, and were amplified under alternating drug conditions. Furthermore, non-genetic resistance was shown to increase the probability that a genetic mutation arises before population extinction in cidal drug environments. Physics-based models have also been proposed to predict AMR [23]. As single-cell approaches are insufficient to capture the dynamics of heterogeneous, interacting cell populations, where resource competition shapes survival and evolutionary outcomes, there is a need to develop population-level models for more realistic biological scenarios [24].

To better understand and mitigate antifungal tolerance and resistance, alternative host models have been developed, particularly using invertebrate animals. Among these, wax moth (*Galleria mellonella*) larvae, have been used to assess the virulence [25], host-pathogens interactions [26], and the efficacy of antifungal treatments [14] against *Candida* and related species. *G. mellonella* larvae are a good infection model as they are inexpensive, easy to maintain in the laboratory, tolerate incubation at human physiological temperatures, and possess cellular and humoral immune responses that share many functional similarities with mammalian innate immunity, though they lack an innate immune system [27, 28]. One important distinction between mammalian and *G. mellonella* innate immunity is the reliance on melanization, a humoral immune response in which melanin is produced around infection sites to trap and kill invading microbes [29]. These characteristics make *G. mellonella* a valuable experimental platform for investigating how infection burden, innate immunity, and antifungal drug treatment collectively shape the dynamics of antifungal tolerance and resistance.

The interplay between drug concentration, tolerance, and resistance has been studied predominantly *in vitro* and in the absence of host immunity [16]. Furthermore, existing treatment strategies tend to focus on drug regimens [30], underestimating the roles of infection burden and host immunity. As a result, it remains unclear how concentration-dependent selection for tolerance versus resistance interacts with host immunity and infection burden to determine treatment outcomes.

We hypothesize that antifungal tolerant and resistant subpopulation dynamics and evolution are affected by drug selection pressure and host immunity. By explicitly integrating immune dynamics with antifungal drug action, this study provides a quantitative framework for predicting the emergence, population dynamics, and evolution of AFR and identifies conditions under which strengthening immune–drug synergy may prove more effective than increasing drug dose alone. These insights are particularly relevant for emerging multidrug-resistant pathogens such as *C. auris* and have direct implications for microbial evolution and clinical treatment.

## 2. Methods

### 2.1. Mathematical model

We develop a deterministic model of the population and evolutionary dynamics of *C. auris* within a host with an innate immune system [31] (e.g., *G. mellonella* larvae). The *C. auris* population is initially represented by drug-susceptible (*S*) and drug-tolerant (*T*) cells, which can reversibly switch between these subpopulations (Fig. 1). T cells can acquire mutations that lead to the evolution of drug-resistant (*R*) cells. These subpopulation transitions were inspired by a previous *in vitro* model [22], which was based on a genetically engineered experimental system [32, 33]. We describe the dynamics of these interacting and evolving subpopulations using a system of coupled ordinary differential equations (ODEs),

**Figure 1.**
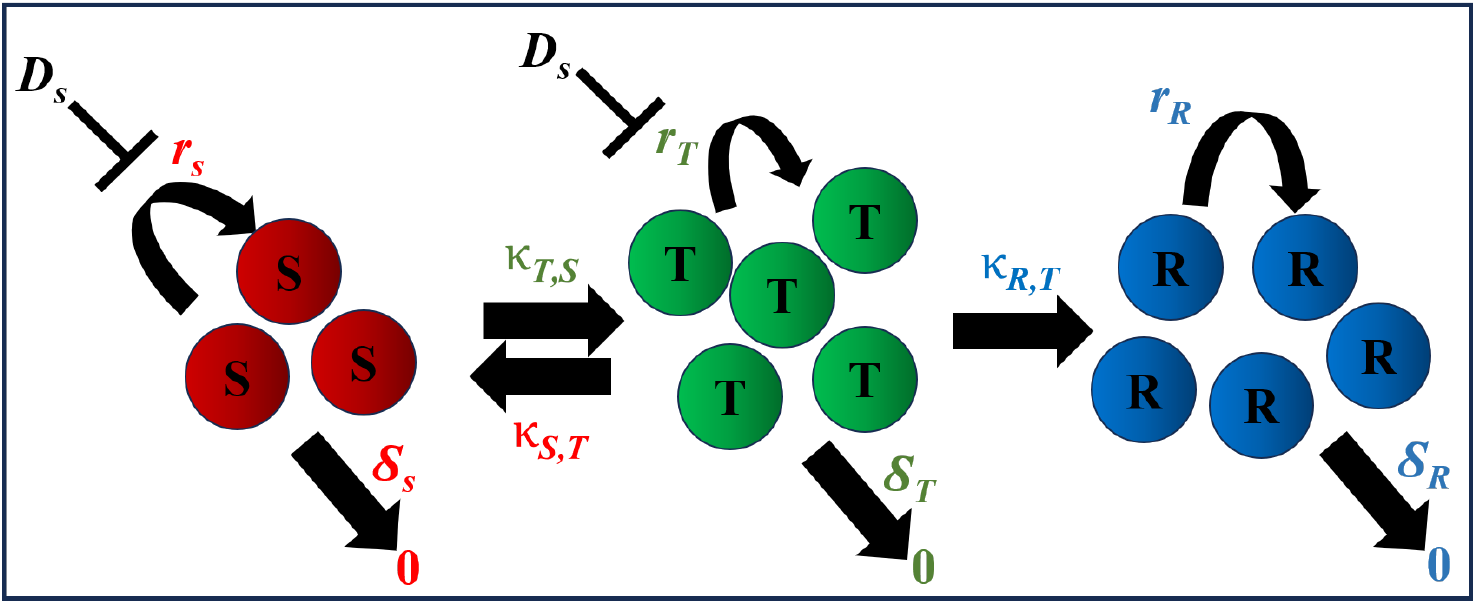
Schematic of our quantitative model depicting the population and evolutionary dynamics of susceptible (S), tolerant (T), and resistant (R) subpopulations for a *C. auris* population under drug treatment and innate immune pressure. Bidirectional arrows represent reversible switching between S and T cells (with rates *κ*_*S,T*_ and *κ*_*T,S*_, respectively), while the unidirectional arrow denotes an irreversible evolutionary transition from T to R cells (with rate *κ*_*R,T*_). Static drugs (represented here by *D*_*S*_) inhibit the growth of the S and T subpopulations. *δ*_*S*_ and *δ*_*T*_ represent the sum effect of cidal drug killing and immune clearance of S and T subpopulations, respectively, and *δ*_*R*_ the immune clearance of the R subpopulation. Transitions are denoted by regular arrows and inhibitions by blunt arrows.

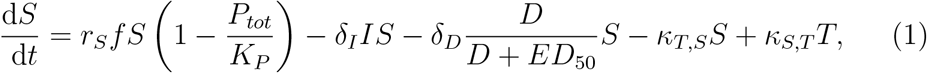

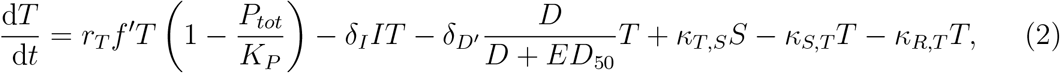

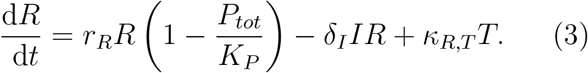

where *r*_*S*_, *r*_*T*_, and *r*_*R*_ are the maximum growth rates of *S, T*, and *R*, respectively. *κ*_*T,S*_ is the switching rate from *S* to *T, κ*_*S,T*_ is the switching rate from *T* to *S*, and *κ*_*R,T*_ is the switching rate from *T* to *R. K*_*p*_ is the maximum cell density. *D* is the antifungal drug concentration and *ED*_50_ is the drug-specific cidal drug concentration that produces a therapeutic effect in 50 % of larvae (Table A1). We assume that a constant number of innate immune cells (*I*) target *C. auris* via a directed search [34]; for simplicity, we do not consider the random search component of a dynamic immune response [35]. Finally, *δ*_*I*_ is the immune clearance rate, and *δ*_*D*_ and *δ*_*D*′_ are the maximum drug-induced killing rates for S and T cells, respectively. We do not model natural basal death rates, as they are assumed to be negligible during antifungal drug treatment.

For static drugs (azoles), the following conditions hold for our ODE model [Eqs. (1)-(3)],

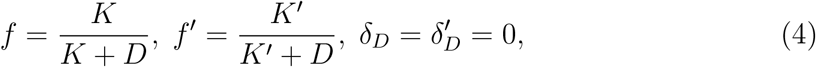

where *f* and *f*′ are functions that describe the growth inhibition of static drugs for *S* and *T* subpopulations, respectively. Correspondingly, *K* and *K*′ are drug-specific half-maximum growth parameters; *K* is estimated from *MIC*_50_ and *K*′ *> K*.

For cidal drugs (polyenes and echinocandins), the following conditions hold for our ODE model [Eqs. (1)-(3)],

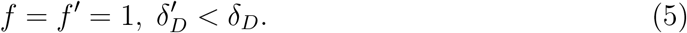

The total population concentration is given by,

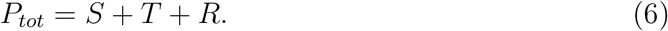

Our population and evolutionary dynamics model assumes density-dependent interactions between the *S, T*, and *R* subpopulations due to competition for a limiting resource; our model does not account for frequency-dependent interactions (e.g., [36, 37]).

### 2.2. Numerical solutions

The system of coupled ODEs [Eqs. (1)-(3)] was solved using MATLAB’s ode45 solver [38]. This solver is well-suited for non-stiff systems and is based on the Dormand–Prince (RKDP) method, an explicit Runge–Kutta method to solve ODEs [39]. The RKDP method estimates the solution by taking a weighted average of slope evaluations at multiple points within each time interval, improving accuracy. RKDP is an adaptive method that dynamically adjusts step sizes to minimize truncation errors while preserving computational efficiency. This makes the RKDP method an ideal choice for quantitatively modeling biological systems with time-dependent dynamics(e.g., [22]).

We define the infection clearance time (*τ*_*clearance*_) as the first time point at which the total *C. auris* population falls below a predefined biologically negligible population density (*P*_*ext*_):

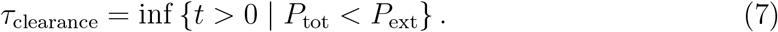

We set *P*_*ext*_ = 1, corresponding to effective *C. auris* eradication within the host. To ensure robustness against transient fluctuations, *τ*_*clearance*_ was only classified as achieved when *P*_*tot*_ *< P*_*ext*_ for the remainder of the simulation. If this condition was not satisfied or if *P*_*tot*_ ≥ *P*_*ext*_ ∀ *t*, then infection clearance was classified as not achieved (*τ*_*clearance*_ = ∞). Clearance times were determined computationally from the numerical ODE solutions.

Simulation time was 200 hours. The parameters for our model are provided in Appendix A (Table A1) and the simulation codes are available at: https://github.com/CharleboisLab/Innate-Immunity.

## 3. Results and Discussion

### 3.1. Susceptible, tolerant, and resistant subpopulation dynamics

In the absence of antifungal treatment, S and T subpopulations rapidly expand to high concentrations under minimal innate immunity and high initial infection load conditions, while the R subpopulation concentration remains negligible (Fig. 2A). Similar growth dynamics occur under minimal innate immunity and low initial infection conditions (Fig. 2C). In contrast, intermediate innate immunity leads to the clearance of the entire *C. auris* population under both low initial infection load (Fig. 2B) and high initial infection load (Fig. 2D), within approximately 20 hours without subsequent regrowth.

**Figure 2.**
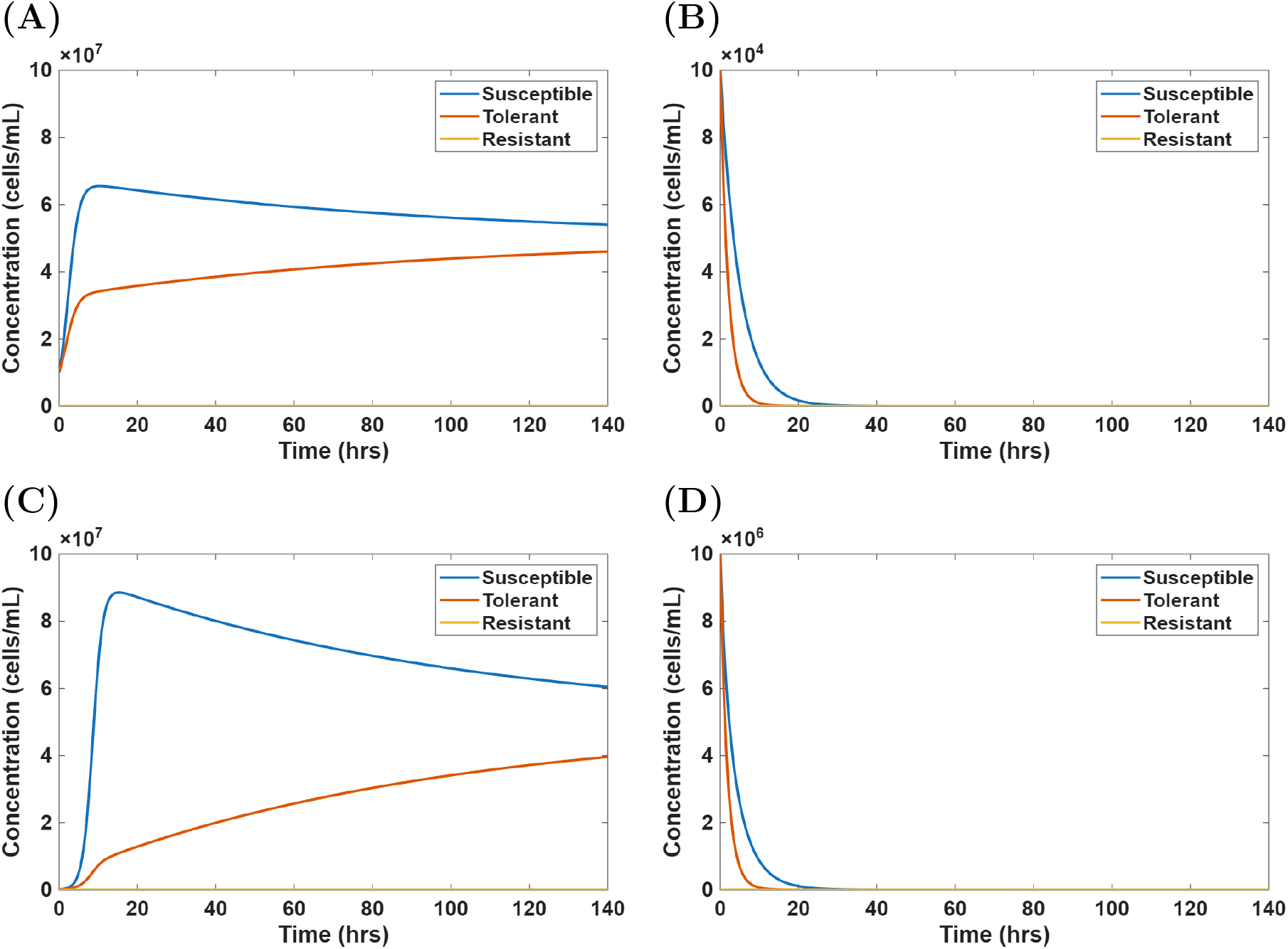
Growth dynamics of a *C. auris* population in the absence of antifungal treatment under different immune and initial infection conditions. **(A)** Minimal innate immunity (10^4^ cells) with a high initial infection load (10^7^ cells/mL). **(B)** Intermediate innate immunity (10^6^ cells) with a low initial infection load (10^5^ cells/mL). **(C)** Minimal innate immunity with a low initial infection load. **(D)** Intermediate innate immunity with a high initial infection load.

For high infection load and intermediate innate host immunity, the S subpopulation rapidly declines during early exposure to both static (Fig. 3A-D) and cidal (Fig. 3E-F) antifungal drugs. Under static azole drug exposure (Fig. 3A-D), a pronounced expansion of the T subpopulation occurred within 20-30 hours, followed by a gradual dominance of the R subpopulation after approximately 30 hours. Under cidal drug exposure with amphotericin B (Figure 3E) and caspofungin (Figure 3F), the T subpopulation reaches higher concentrations more quickly and dominates the population until around 80 hours, after which the T subpopulation declines; after about 100 hours, the R subpopulation dominates the population. In contrast, simulations incorporating strong innate immunity quickly lead to total population collapse (data not shown).

**Figure 3.**
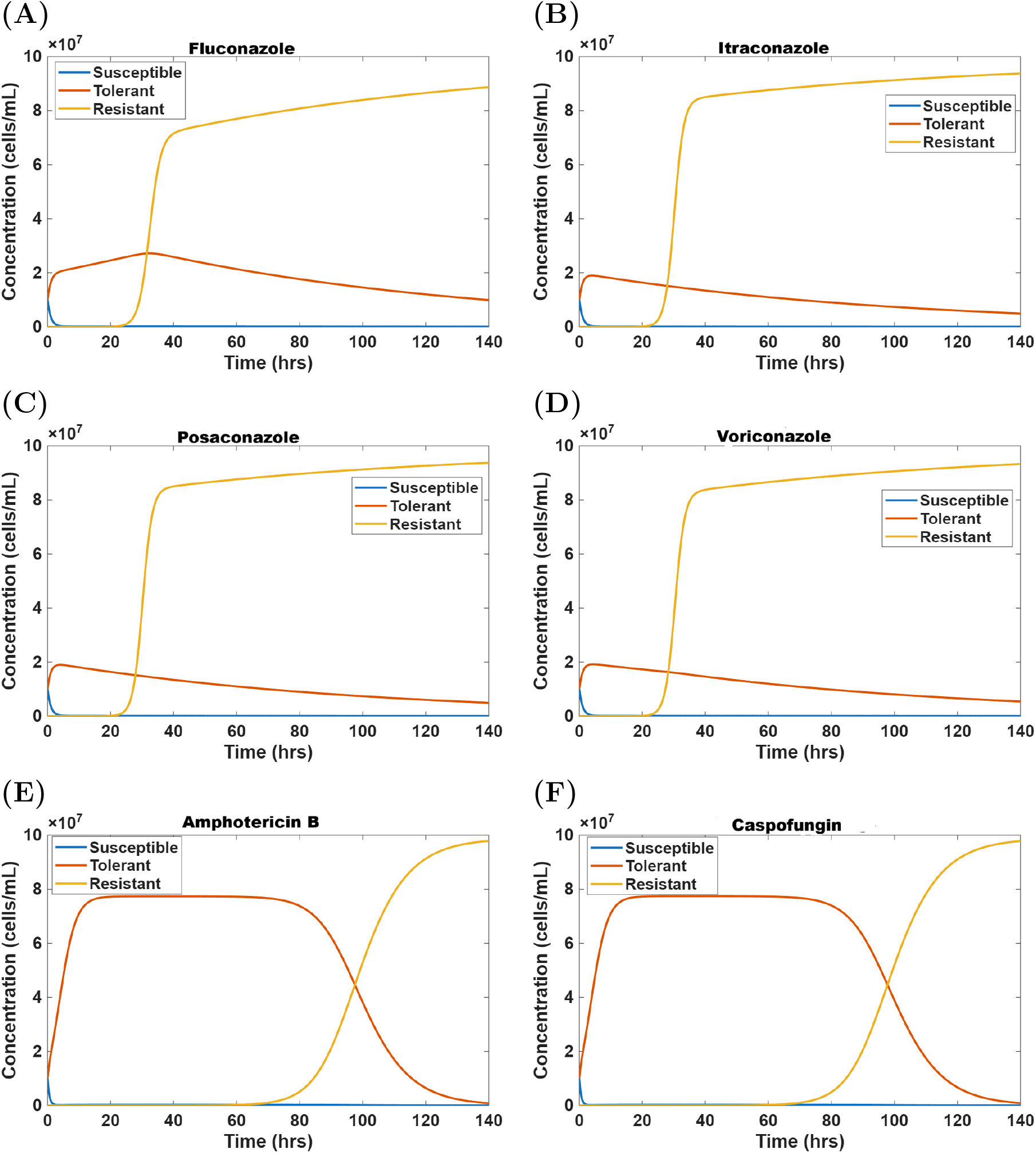
Growth dynamics of a *C. auris* population under antifungal treatment in intermediate innate immunity and high infection load conditions. Simulations show the subpopulation trajectories of susceptible, tolerant, and resistant subpopulations during exposure to **(A)** fluconazole, **(B)** itraconazole, **(C)** posaconazole, **(D)** voriconazole, **(E)** amphotericin B, and **(F)** caspofungin antifungal drugs. All simulations were performed at a drug concentration of 640 *µ*g/mL with an intermediate innate immunity (10^6^ cells) and a high initial infection load (10^7^ cells/mL).

These simulations predict that acute antifungal tolerance allows *C. auris* populations to survive static and cidal antifungal drugs until permanent resistance evolves (Fig. 3). This phenomenon is in agreement with previous work on the transition from non-genetic to genetic drug resistance [22, 20]. However, this previous modeling work did not incorporate innate immunity and found that resource competition between non-genetically and genetically resistant subpopulations always hindered resistance evolution for static and cidal drug treatments. We find that when innate host immunity is incorporated into the model, resource competition between T and R subpopulations delays the evolution of resistance for cidal drugs, but not for static drugs (Fig. 3). We note that although non-genetically resistant and tolerant phenotypes can both revert to the susceptible phenotype [15, 22], the molecular mechanisms underlying antifungal tolerance are still being elucidated and may not be non-genetic in nature [15, 12].

### 3.2. Tolerance and resistance crossover times

Next, we investigate how drug class and concentration, infection burden, and the innate immune response affect the crossover time at which the T and R subpopulations exchange population dominance (Figs. 4-6).

**Figure 4.**
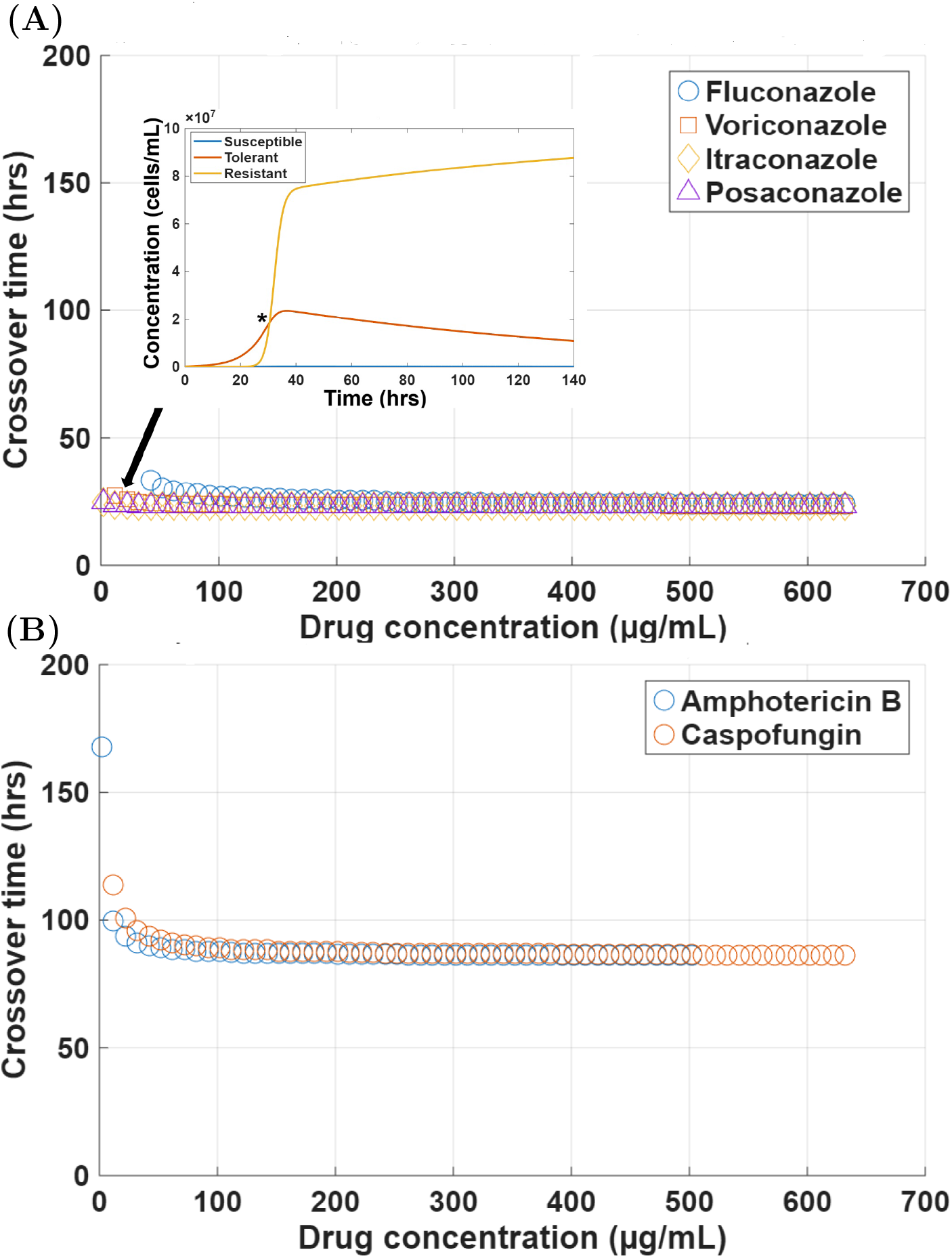
Crossover times between tolerant and resistant subpopulations for minimum innate immunity and a low initial infection under antifungal drug treatment. **(A)** Static antifungal treatment (fluconazole, voriconazole, itraconazole, and posaconazole). The inset shows the crossover event (indicated by the asterisk) under static drug action at a low concentration (6 *µ*g/mL) for voriconazole. **(B)** Cidal antifungal treatment (amphotericin B and caspofungin). Minimum innate immunity corresponds to 10^4^ cells and low initial infection load to 10^5^ cells/mL.

For minimum innate immunity and low initial infection load, the crossover times are largely independent of drug concentration (Fig. 4). For azole antifungal drugs the crossover events occur within a narrow temporal window of approximately 20 to 30 hours (Fig. 4A). A notable exception is that cross over events do not occur during our simulations for fluconazole below approximately 40 *µ*g/ml. This is either due to the population dynamics, such that R cannot dominate T in the population, or the simulation timescale (Sec. 2.2) is not sufficiently long to observe them. The crossover times for the cidal antifungal drugs depend more strongly on low drug concentrations (Fig. 4B). In the low concentration regime, the crossover times exceed 100 hours for caspofungin and 150 hours for amphotericin B. As drug concentration increases, crossover times decrease rapidly and plateau at around 85 hours. The crossover time profiles for static and cidal antifungal drugs for minimum innate immunity and intermediate initial infection load were qualitatively similar (Fig. B1 in Appendix B) to those of minimum innate immunity and low initial infection load (Fig. 4).

At a minimum innate immunity and high initial infection load (Fig. 5), the sensitivity of crossover time events to static antifungal drugs becomes more pronounced (Fig. 5A). Fluconazole shows an exponential decline in the crossover time over a large concentration range, with crossover events occurring beyond 100 hours at low to intermediate concentrations and converging to approximately 30 hours at high concentrations. Voriconazole, itraconazole, and posaconazole exhibit similar dynamics at lower concentrations. In contrast, cidal antifungal drugs have concentration-crossover time profiles similar to those of low (Fig. 4) and intermediate initial infection loads (Fig. B1 in Appendix B).

**Figure 5.**
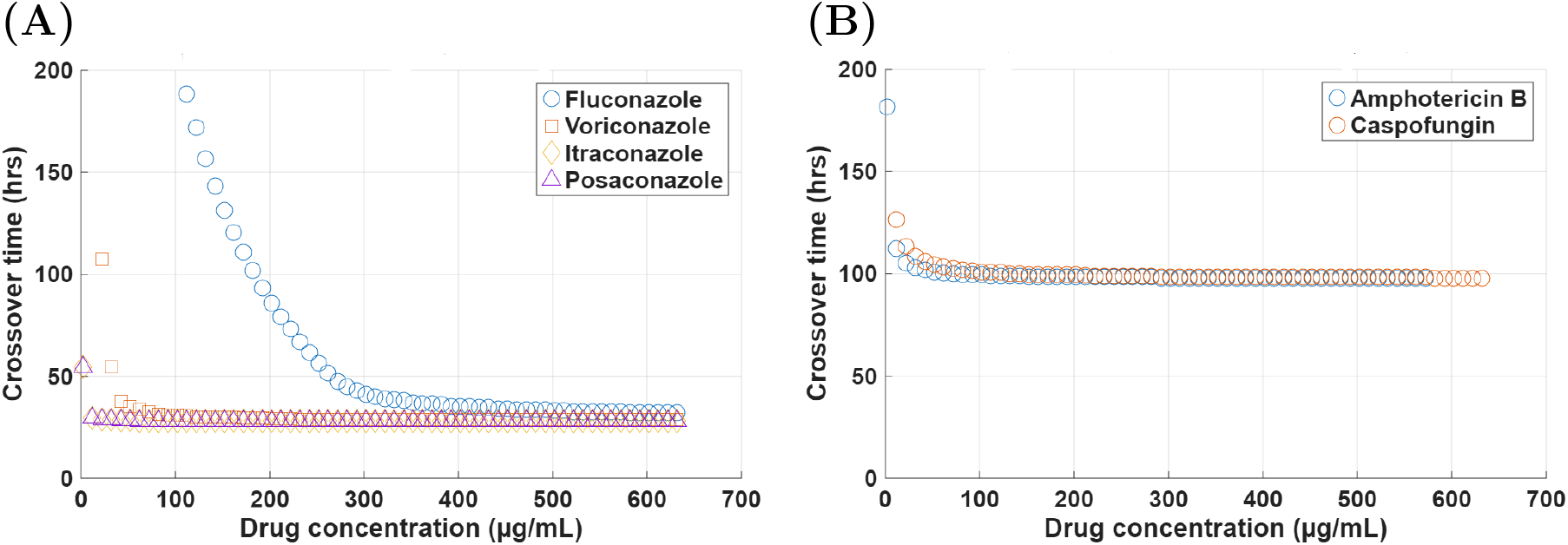
Crossover times between tolerant and resistant subpopulations for minimum innate immunity and a high initial infection under antifungal drug treatment. **(A)** Static antifungal treatment (fluconazole, voriconazole, itraconazole, posaconazole). **(B)** Cidal antifungal treatment (amphotericin B and caspofungin). Minimum innate immunity corresponds to 10^4^ cells and high initial infection load to 10^7^ cells/mL.

When the host has an intermediate level of innate immunity and low initial infection, T and R crossover times are constant across the concentrations of all antifungal drugs (Fig. 6). For static drug treatments, crossover times are close to zero (approximately 3 hours; Fig. 6A). For cidal drug treatments, crossover times are consistently above zero (approximately 45 *µ*g/ml) across drug concentrations (Fig. 6B). For intermediate innate immunity with intermediate infection, the azoles at low drug concentration (less than 50ug/ml), crossover times are within 15-40 hrs, then stabilize to 15-30 hrs after drug concentrations greater than 50 ug/ml (Fig. B2A in Appendix B). For cidal drugs, the crossover time profiles are similar to those for intermediate immunity and low infection (Fig. 6B).

**Figure 6.**
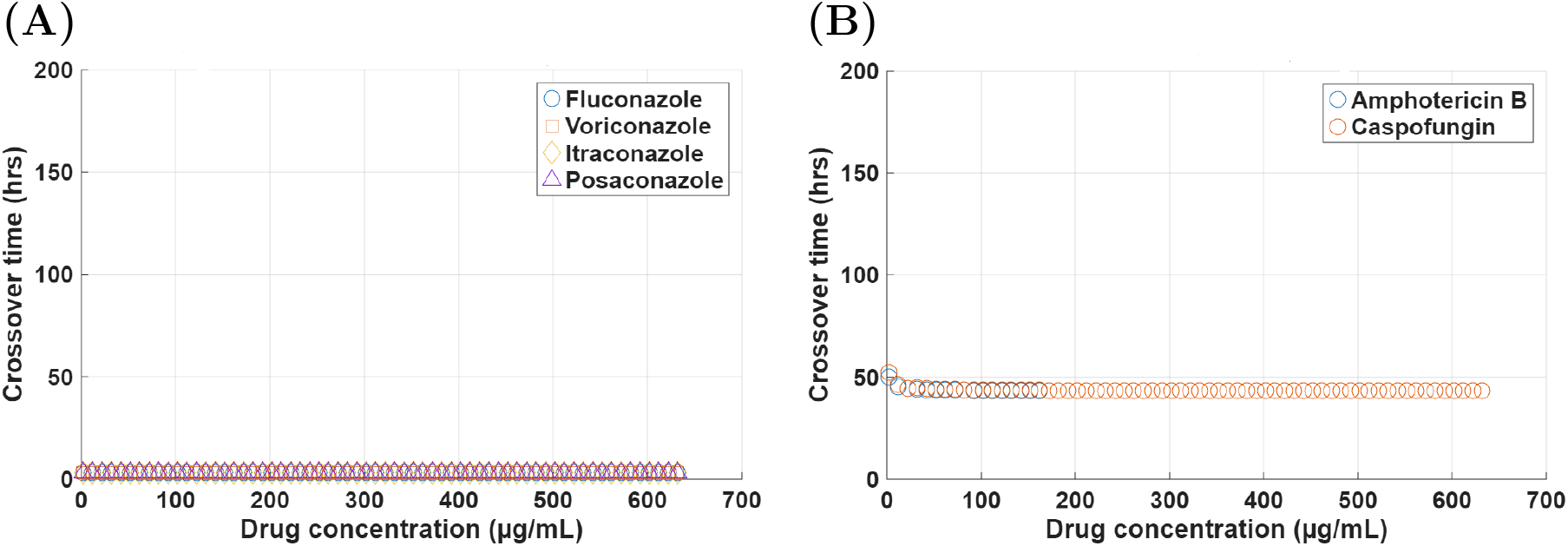
Crossover times between tolerant and resistant subpopulations for intermediate innate immunity and a low initial infection under antifungal drug treatment. **(A)** Static antifungal treatment (fluconazole, voriconazole, itraconazole, posaconazole). **(B)** Cidal antifungal treatment (amphotericin B and caspofungin). Intermediate innate immunity corresponds to 10^6^ cells and minimal initial infection load to 10^5^ cells/mL.

For intermediate innate immunity, high initial infection, the crossover times for static drugs are within 20 to 40 hours at low drug concentration (less than 50 *µ*g/ml) and then stabilize after 20 to 25 hours at 50 *µ*g/ml (Fig. B2B in Appendix B). For cidal drugs, the crossover time profiles are similar to those for intermediate immunity and low infection (Fig. 6B).

As crossover times saturate at high drug concentrations, further dose increases offer diminishing returns for modulating population dynamics and evolutionary outcomes. Subpopulation crossovers do not occur for maximum innate immunity, as the T and R subpopulations are cleared from the host before they can establish in the population (data not shown).

These findings reveal that the crossover times between T and R subpopulations are governed by the interplay between antifungal drug class and concentration, infection burden, and the host’s innate immune system. This highlights that optimal antifungal treatment strategies should consider these factors, as they may be critical to prevent the emergence of antifungal tolerance and resistance.

### 3.3. Infection clearance times

For antifungal treatment with static drugs, S and T subpopulations clear rapidly once innate immunity levels exceed 10^5^ cells (Fig. 7A). Below this threshold, clearance times typically exceed 100 hours, reflecting ineffective infection control of a weak immune system even during drug treatment. At lower levels of immunity, the T subpopulation survives slightly longer compared to the S subpopulation during static drug treatment due to antifungal tolerance. The R subpopulation remains uncleared (*τ*_*clearance*_ = ∞) throughout the innate immunity range (data not shown), despite drug treatment and increasing immunity levels, due to antifungal drug resistance. Similar infection clearance dynamics occur for antifungal treatment with cidal drugs (Fig. 7B).

**Figure 7.**
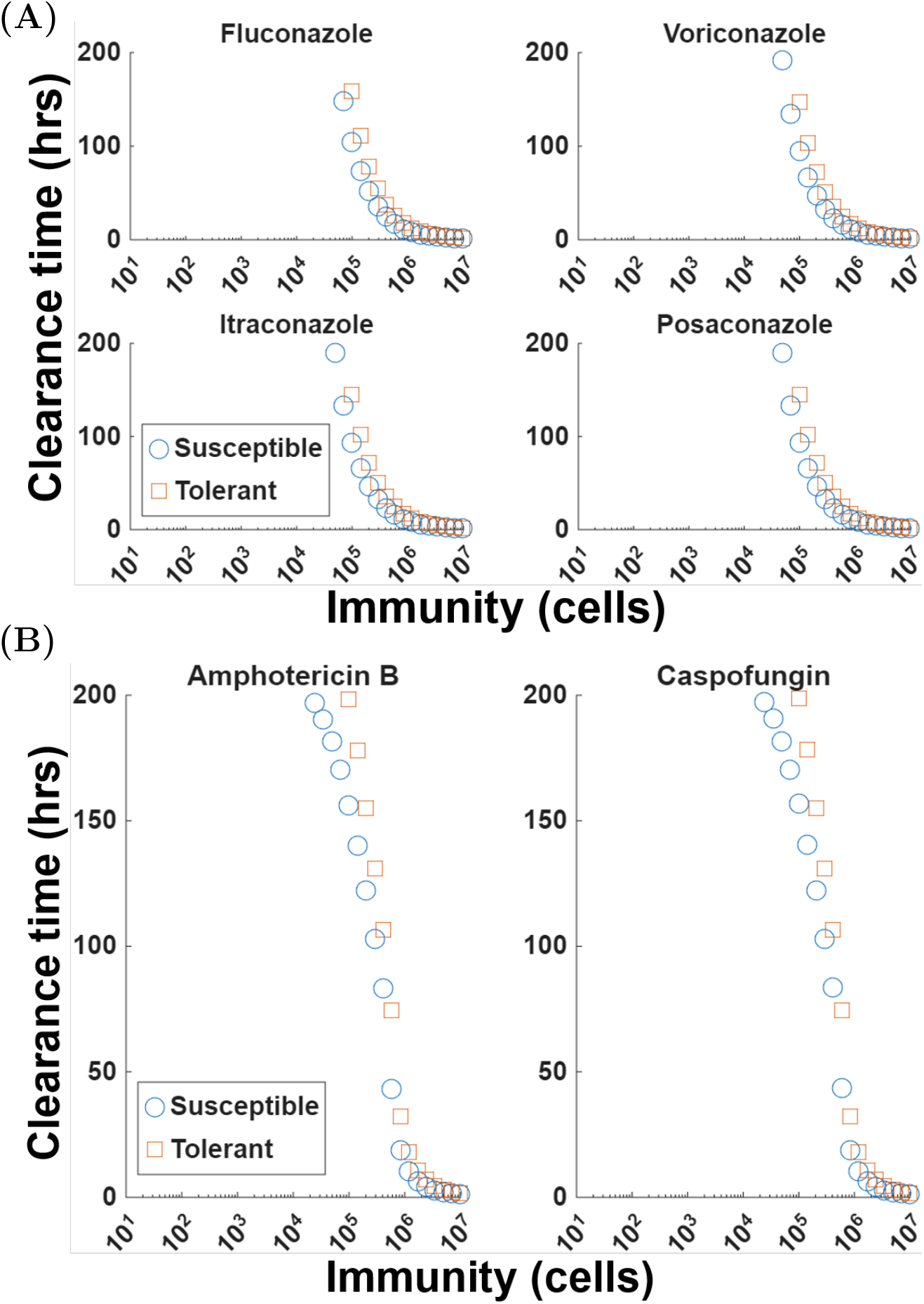
Innate immunity-dependent clearance of *C. auris* susceptible, tolerant, and resistant subpopulations for antifungal treatment of an intermediate initial infection. High concentration (640 *µ*g/mL) **(A)** static drug and **(B)** cidal drug treatments. Intermediate initial infection load corresponds to 10^6^ cells/mL. Resistant subpopulations are not shown, as they are not cleared during the simulation timeframe.

These results highlight the critical role of host immunity in determining infection clearance during antifungal therapy. Antifungal drugs alone, whether static or cidal, may be insufficient to eliminate tolerance and resistance, underscoring the importance of immunocompetence in *C. auris* infection control [40, 14, 41].

### 3.4. Tolerance versus resistance regimes

We found that antifungal tolerance can emerge as the dominant subpopulation early during drug exposure. For the low infection condition, the T subpopulation dominates the population during fluconazole (Fig. 8A) and caspofungin (Fig. 8B) treatments. Similarly, the T subpopulation dominates the intermediate infection condition for fluconazole, voriconazole, and caspofungin treatments (Fig. 9A-C, respectively) and the high infection condition for fluconazole, itraconazole, voriconazole, and caspofungin treatments (Fig. 10A-D, respectively). The S and R subpopulations do not establish in the population in any of these scenarios beyond near-zero or low concentrations, except for the R subpopulation, which establishes at a similar concentration to the T subpopulation when exposed to itraconazole during high infection (Fig. 10B).

**Figure 8.**
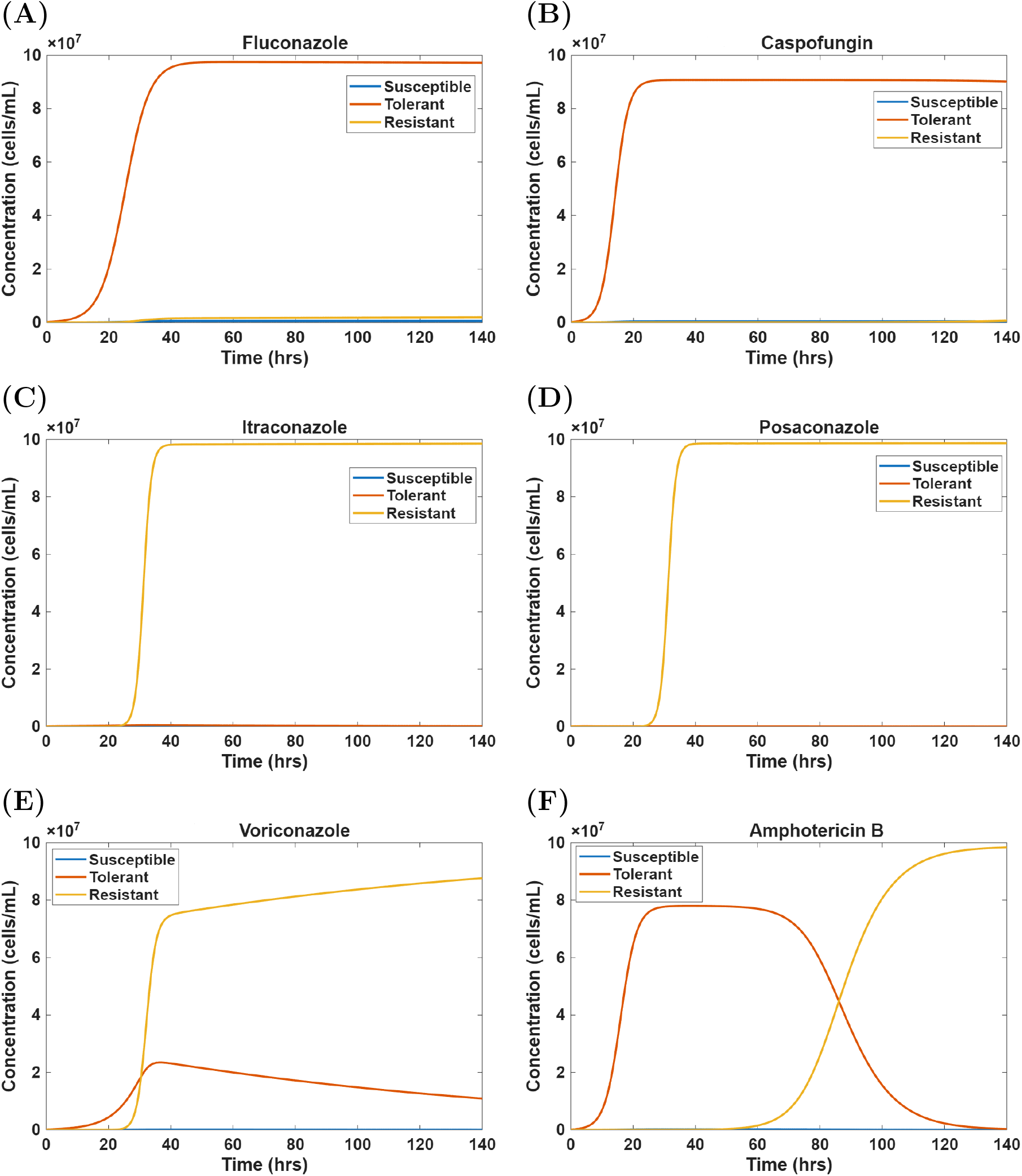
Competition between susceptible, tolerant, and resistant *C. auris* subpopulations during low initial infection and minimum innate immunity. The tolerant subpopulation dominates the population during treatment with **(A)** fluconazole (12 *µ*g/mL) and **(B)** caspofungin (2 *µ*g/mL). The resistant subpopulation emerges as the dominant subpopulation during treatment with **(C)** itraconazole (3 *µ*g/mL), **(D)** posaconazole (40 *µ*g/mL), **(E)** voriconazole (6 *µ*g/mL), and **(F)** amphotericin B (50 *µ*g/mL). Low initial infection load corresponds to 10^5^ cells/mL and minimum innate immunity corresponds to 10^4^ cells.

**Figure 9.**
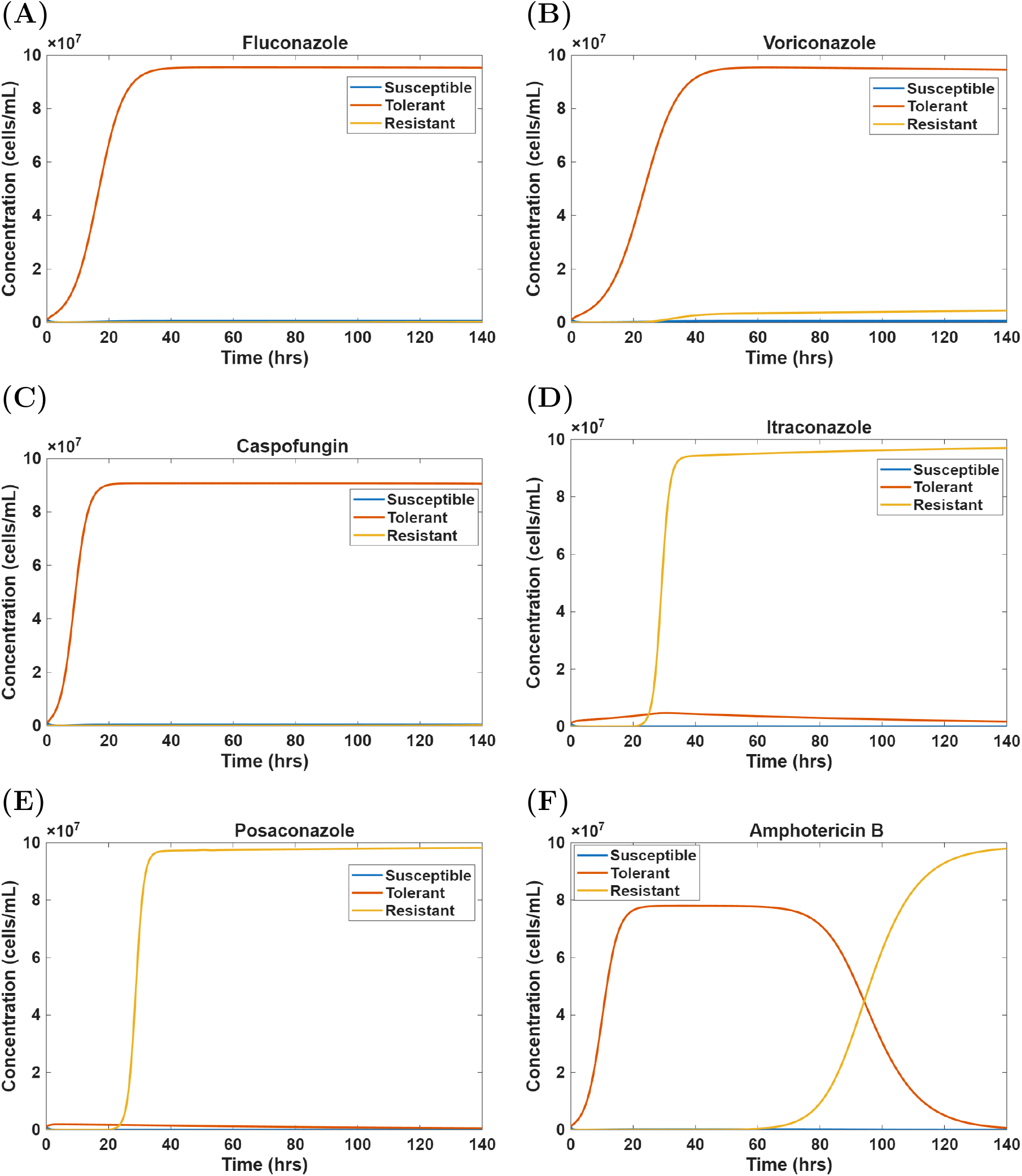
Competition between susceptible, tolerant, and resistant *C. auris* subpopulations during intermediate initial infection and minimum innate immunity. The tolerant subpopulation dominates the population during treatment with **(A)** fluconazole (12 *µ*g/mL), **(B)** voriconazole (6 *µ*g/mL), and **(C)** caspofungin (2 *µ*g/mL). The resistant subpopulation emerges as the dominant subpopulation during treatment with **(D)** itraconazole (3 *µ*g/mL), **(E)** posaconazole (40 *µ*g/mL), and **(F)** amphotericin B (50 *µ*g/*mL*). Intermediate initial infection load corresponds to 10^6^ cells/mL and minimum innate immunity corresponds to 10^4^ cells.

**Figure 10.**
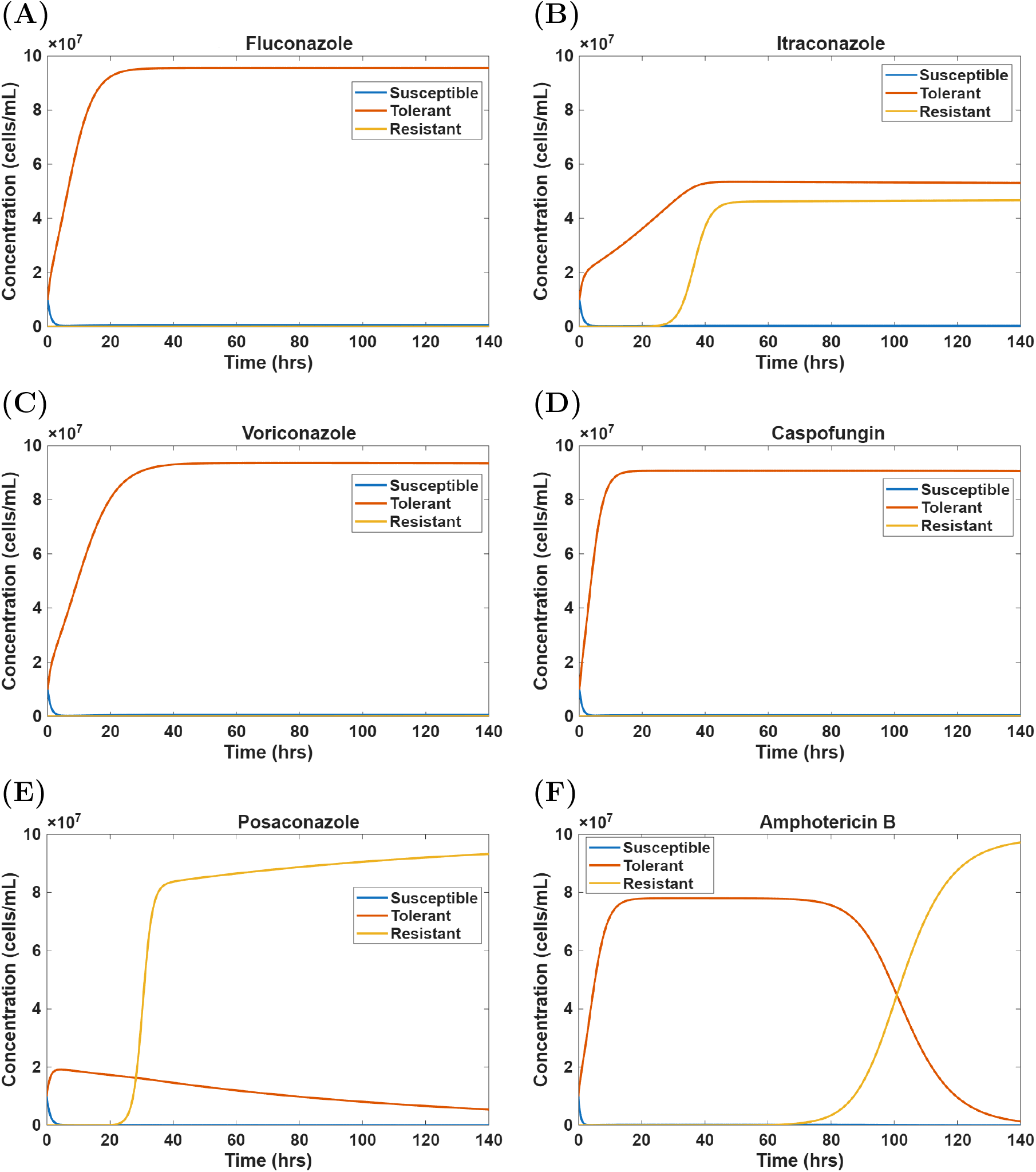
Competition between susceptible, tolerant, and resistant *C. auris* subpopulations during high initial infection and minimum innate immunity. The tolerant subpopulation dominates the population during treatment with **(A)** fluconazole (12 *µ*g/mL), **(B)** itraconazole (3 *µ*g/mL), **(C)** voriconazole (6 *µ*g/mL), and **(D)** caspofungin (2 *µ*g/mL). The resistant subpopulation emerges as the dominant subpopulation during treatment with **(E)** posaconazole (40 *µ*g/mL) and **(F)** amphotericin B (50 *µ*g/mL). High initial infection load corresponds to 10^7^ cells/mL and minimum innate immunity corresponds to 10^4^ cells.

R emerges as the dominant subpopulation in the low infection condition during itraconazole, posaconazole, voriconazole, and amphotericin B treatments (Fig. 8C-F, respectively). R also emerges as the dominant subpopulation in the intermediate infection condition during treatment with itraconazole, posaconazole, and amphotericin B (Fig. 9D-F, respectively) and in the high infection condition during posaconazole and amphotericin B treatments (Fig. 10E,F). T and R crossover times vary across drug, infection, and immunity conditions (Sec. 3.2).

Under conditions of minimum innate immunity, either the T or R subpopulation emerges as dominant, depending on if the antifungal drug is static or cidal. For fungistatic drugs, this behavior is primarily determined by the parameter *K*′, which was set to the corresponding *MIC*_50_ value for each drug against *C. auris* (Table A1 in Appendix A). This parameter modulates fungal growth during drug exposure [Eqs. (1)-(2)]. The T subpopulation dominates in the presence of fluconazole (Fig. 8A, Fig. 9A, and Fig. 10A) because fluconazole has the highest associated *K*′ value among the antifungal drugs considered. Similarly, T dominates under voriconazole treatment (Fig. 9B and Fig. 10C), which has the second-highest *K*′ value. In contrast, for itraconazole, the T subpopulation dominates only at high infection burdens and only during the early stages of infection (Fig. 10B). However, after approximately 850 hours, the R subpopulation overtakes the T subpopulation (Fig. B3 in Appendix B). Note that in our model *K*′ for T is greater than *K* for S (Sec. 2.1), which explains why T dominates over S in these scenarios.

For fungicidal drugs, the dominant subpopulation is primarily determined by the *ED*_50_ parameter (Table A1), which governs drug efficacy [Eqs. (1)-(2)]. The T subpopulation dominates under caspofungin treatment (Fig. 8B, Fig. 9C, and Fig. 10D) because caspofungin has a lower *ED*_50_ value than amphotericin B. A lower *ED*_50_ indicates greater drug potency, resulting in a stronger cidal effect at a given drug concentration. This allows T subpopulation to rapidly outcompete the S subpopulation, because the drug-induced killing rate of the T subpopulation (*δ*_*D*′_) is lower than for the S subpopulation (*δ*_*D*_) for the same *ED*_50_. Due to the cidal effect of the drug on T cells, the R subpopulation is initially deprived of a large T subpopulation from which to mutate. As *T* → *K*_*p*_, the T subpopulation becomes large enough to act as a substantial source of R cells, but in this regime the T subpopulation dominates the R subpopulation by hoarding the shared resources [i.e., *P*_*tot*_ in Eq. (6) is mainly comprised of *T* cells, which suppresses the growth of the R subpopulation].

Overall, the T subpopulation generally emerges as the dominant subpopulation or survives long enough to act as a substrate for the evolution of the antifungal-resistant R subpopulation. The latter phenomenon is known for the evolution of genetic resistance from non-genetically resistant subpopulations [22, 20, 21], but not from tolerance. The former phenomenon, to our knowledge, is a novel prediction. The T and R subpopulations can also co-exist in our simulations for some infection, drug, and immunity conditions (e.g., Fig. 10B).

## 4. Conclusion

We developed a quantitative, population-level model based on the experimental *C. auris*-*G. mellonella* infection model, including the incorporation of experimentally-derived infection burden, immune strength, fungal growth, and antifungal drug parameters. By explicitly incorporating experimentally measurable quantities of the larval host, such as fungal burden and hemocyte-mediated immune responses, with population-level evolutionary dynamics, our model provides a biologically grounded framework to gain insights into and predict antifungal tolerance and resistance *in vivo*.

Our study predicted that *C. auris* susceptible, tolerant, and resistant subpopulation dynamics in *G. mellonella* larvae are governed by infection burden, immune-mediated clearance, and antifungal drugs. When incorporating these biologically and clinically relevant factors into our model, we uncovered previously unobserved phenomena, including the establishment and dominance of tolerant subpopulations over resistant subpopulations, as well as the co-existence of these subpopulations. Previous work, which did not consider immune responses, found that non-genetic resistance can facilitate survival [20, 21, 42] while hindering the evolution of genetic drug resistance [22].

The modeling framework developed in this study is deterministic, which provides an accurate and computationally efficient description of microbial populations at high densities. However, the deterministic nature of our model limits its applicability during the initial infection period or when antifungal drug treatment and/or immune clearance have reduced the infecting microbes to low levels, where stochastic effects may be important for population dynamics and evolution [24]. Accordingly, subsequent modeling should include stochastic simulations to investigate the emergence of tolerance and resistance in low concentration regimes (e.g., [22]). We also focused on innate immunity, which provides a rapid, generalized, non-specific defense against invading pathogens, whereas the adaptive immune system, which provides a slower, specialized, and specific defense, is not considered [43, 44]. Incorporating time-delayed immune activation through delay stochastic simulation approaches [45, 46] in the model would provide a more realistic representation of host-pathogen interactions than the instantaneous innate immunity that we considered. Finally, our model should be expanded to incorporate the adaptive immune system for relevant host organisms.

The modeling predictions in this study can be tested *in vivo* by varying the infection burden [25, 47], immune strength [26, 48], and antifungal treatment [14, 49] in *G. mellonella* larvae infected with *C. auris*. Qualitatively, the presence of antifungal susceptible, tolerant, and resistant subpopulations can be identified via established *in vitro* assays [15, 50, 19] from cultures obtained from larval hemolymph. Experimental measurements of susceptible, tolerant, and resistant subpopulations can be performed and compared to our predicted crossover and clearance times. This could be done by survival assays, fungal burden quantification, and time-kill analyses of extracted larval hemolymph [27, 51, 52]. Fluctuation analyses [53, 54] could estimate the mutation rates associated with transitions from tolerance to resistance under antifungal drug pressure. Molecular techniques can identify common antifungal resistance mutations/genes [55]. However, quantitatively determining the dynamics of infecting subpopulations in a host organism will be challenging, in part because the molecular mechanisms underlying tolerance in *C. auris* and related *Candida* species have yet to be fully elucidated [15]. Though, genetic signatures, including aneuploidy, may be used to quantify tolerant subpopulations [12].

Overall, we provided a quantitative framework for investigating infecting pathogen populations under immune and drug selection pressures. Our findings expand our understanding of the interactions between different forms of antimicrobial resistance. This study moves us one step closer to the ultimate goal of developing a quantitative and predictive understanding of antimicrobial resistance [23] and using this knowledge to guide the treatment of patients with life-threatening infections.

## Author Contributions

D.A.C. conceived and designed the study. D.A.C. and C.M.G. developed the mathematical models. C.M.G. performed the numerical simulations and visualized the data. C.M.G. and D.A.C. analyzed the data. C.M.G. and D.A.C. wrote the article. D.A.C. supervised and obtained funding for the research.

## Acknowledgments

D.A.C. was financially supported by the Human Frontier Science Program (RGEC30/2024) and by the Natural Sciences and Engineering Research Council of Canada (RGPIN-2020-04007). C.M.G. was supported by the DOST-SEI-UAlberta S&T Graduate Scholarship program.

## Data availability statement

Data supporting the findings of this study are openly available at the following URL/DOI: https://github.com/CharleboisLab/Innate-Immunity.

## Appendix A. Parameters

The parameter values used in our study (Table A1) were based on experimental studies on *C. auris*. The initial susceptible and tolerant subpopulation concentrations *S*_0_ and *T*_0_, respectively, were informed by ongoing wax moth (*Galleria mellonella*) larvae-*C. auris* infection experiments in the Charlebois Laboratory. The protocols for these experiments are described in the literature [56, 57, 27, 58]. The resistant subpopulation *R*_0_ concentration was set to zero, as we are modeling *C. auris* strains that are initially susceptible/tolerant (not initially resistant) to antifungal drugs. The growth rate parameters *r*_*S*_, *r*_*T*_, and *r*_*R*_, were assigned two sets of values to capture different treatment conditions. In particular, the case where *r*_*R*_ = 0 corresponds to the no-drug scenario in Figure 2, where the resistant subpopulation is absent. Under this no-drug condition, *r*_*S*_ = 0.8 was used to reflect the higher growth of the susceptible subpopulation. Drug concentrations (*D*) were selected based on the Infectious Diseases Society of America Clinical Practice Guidelines for human antifungal therapy [59]. For our simulations, these concentrations were scaled to ten times the doses of the clinical antifungal therapy, as was done in *G. mellonella* larvae infection experiments [60].

## Appendix B. Supporting Simulations

The crossover times for an intermediate initial infection load are shown in Figure B1. These crossover times are qualitatively similar to those for a low initial infection load (Fig. 4). Fluconazole exhibits a pronounced concentration-dependent decrease in the crossover time from low to intermediate doses, followed by a convergence of the crossover time to those of the other drugs (around 25 *µ*g/ml) at higher concentrations. Cidal drugs again have less concentration dependent crossover times. Crossover times decrease rapidly as amphotericin B and caspofungin concentrations increase, and then plateau at approximately 90 hours (Fig. B1).

**Table A1:** Parameters used for numerically simulating the deterministic *in vivo* evolutionary drug resistance dynamics in static and cidal drug conditions. A horizontal line in the ‘Cidal Drug Value’ column indicates that the value is the same as the corresponding value in the ‘Static Drug Value’ column.

| Parameter | Static Drugs | Cidal Drugs | Units | Reference |
| --- | --- | --- | --- | --- |
| $r_S$ | 0, 0.8 | - | $h^{-1}$ | [22] |
| $r_T$ | 0.3, 0.5 | - | $h^{-1}$ | [22, 61, 62] |
| $r_R$ | 0.8, 0 | - | $h^{-1}$ | [61, 62] |
| $S_0$ | $10^5 - 10^7$ | - | $cells\ mL^{-1}$ | - |
| $T_0$ | $10^5 - 10^7$ | - | $cells\ mL^{-1}$ | - |
| $R_0$ | 0 | - | $cells\ mL^{-1}$ | - |
| $K_P$ | $10^8$ | - | $cells\ mL^{-1}$ | [63, 64] |
| $\delta_I$ | $10^{-6}$ | - | $h^{-1}$ | [65, 62] |
| $I$ | $10^1 - 10^7$ | - | # of hemocytes | [31, 66, 62] |
| $\delta_D$ | 0 | 1.0 | $h^{-1}$ | [22] |
| $\delta_{D'}$ | 0 | 0.1 | $h^{-1}$ | [22] |
| $D$ | 2 - 640 | - | $\mu g\ mL^{-1}$ | [59, 15, 60] |
| $K_{flu}$ | 2 | 0 | $\mu g\ mL^{-1}$ | [15] |
| $K_{vor}$ | 0.25 | 0 | $\mu g\ mL^{-1}$ | [15] |
| $K_{itra}$ | 0.25 | 0 | $\mu g\ mL^{-1}$ | [15] |
| $K_{pos}$ | 0.25 | 0 | $\mu g\ mL^{-1}$ | [15] |
| $K'_{flu}$ | 64 | 0 | $\mu g\ mL^{-1}$ | [15] |
| $K'_{vor}$ | 8 | 0 | $\mu g\ mL^{-1}$ | [15] |
| $K'_{itra}$ | 0.5 | 0 | $\mu g\ mL^{-1}$ | [15] |
| $K'_{pos}$ | 0.5 | 0 | $\mu g\ mL^{-1}$ | [15] |
| $ED_{50, azoles}$ | 0 | 0 | $\mu g\ mL^{-1}$ | [15] |
| $ED_{50, amb}$ | 0 | 2 | $\mu g\ mL^{-1}$ | [15] |
| $ED_{50, cas}$ | 0 | 4 | $\mu g\ mL^{-1}$ | [15] |
| $\kappa_{T,S}$ | $9.7 \times 10^{-1}$ | - | $h^{-1}$ | [67] |
| $\kappa_{S,T}$ | $5.3 \times 10^{-3}$ | - | $h^{-1}$ | [67] |
| $\kappa_{R,T}$ | $10^{-8}$ | - | $h^{-1}$ | [68, 69, 62] |

**Figure B1:**
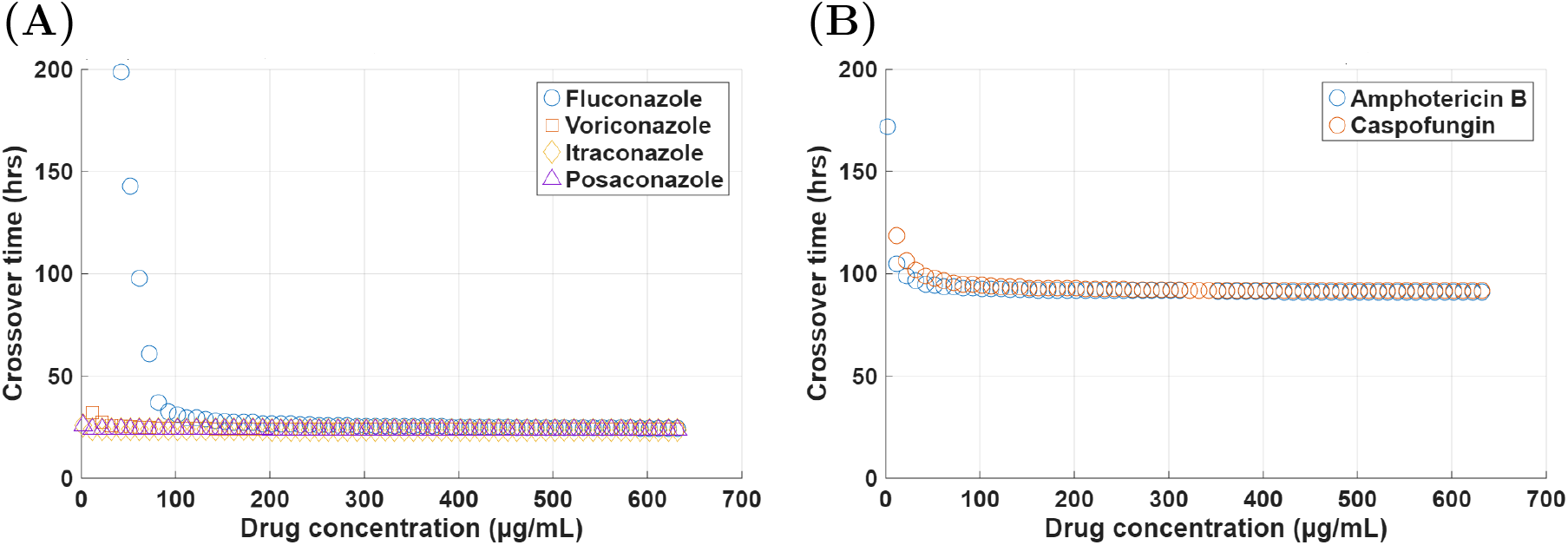
Crossover times between tolerant and resistant subpopulations for minimum innate immunity and an intermediate initial infection under antifungal drug treatment. **(A)** Static antifungal treatment (fluconazole, voriconazole, itraconazole, and posaconazole). **(B)** Cidal antifungal treatment (amphotericin B and caspofungin). Minimum innate immunity corresponds to 10^4^ cells and intermediate initial infection load to 10^6^ cells/mL.

**Figure B2:**
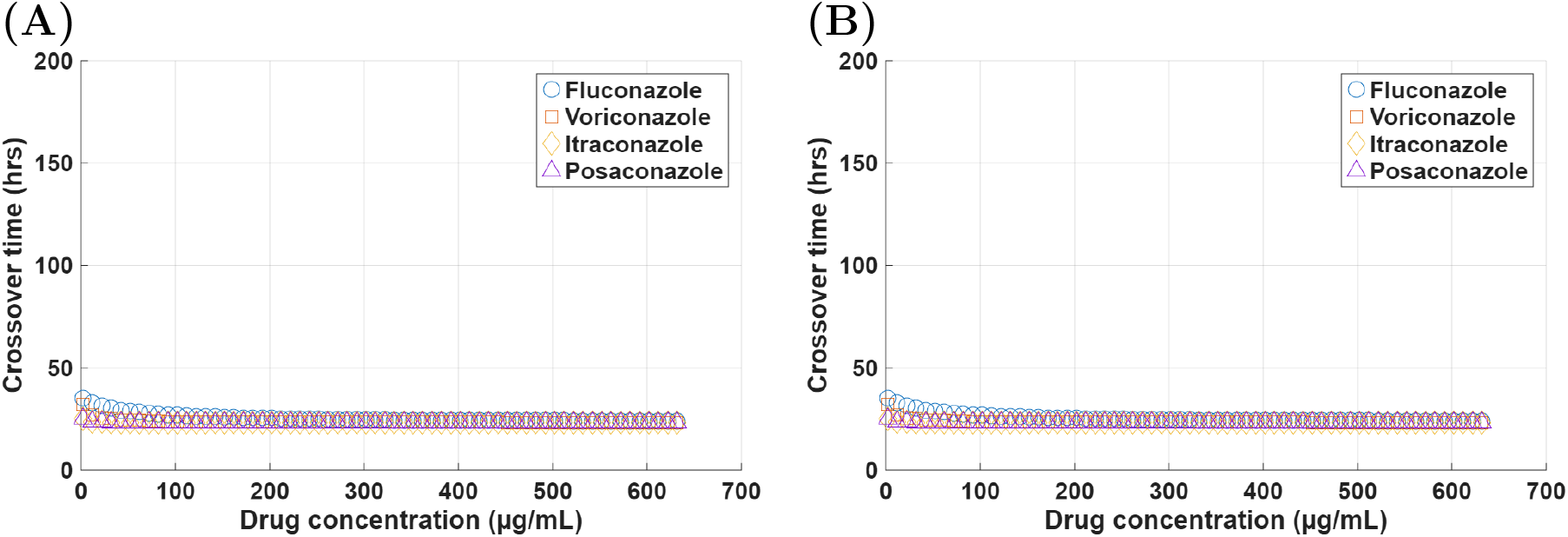
Crossover times between tolerant and resistant subpopulations for **(A)** intermediate innate immunity and an intermediate initial infection and **(B)** intermediate innate immunity and a high initial infection under static antifungal treatment (fluconazole, voriconazole, itraconazole, posaconazole). Intermediate innate immunity corresponds to 10^6^ cells, intermediate initial infection load to 10^6^ cells/mL, and high initial infection load to 10^7^ cells/mL.

**Figure B3:**
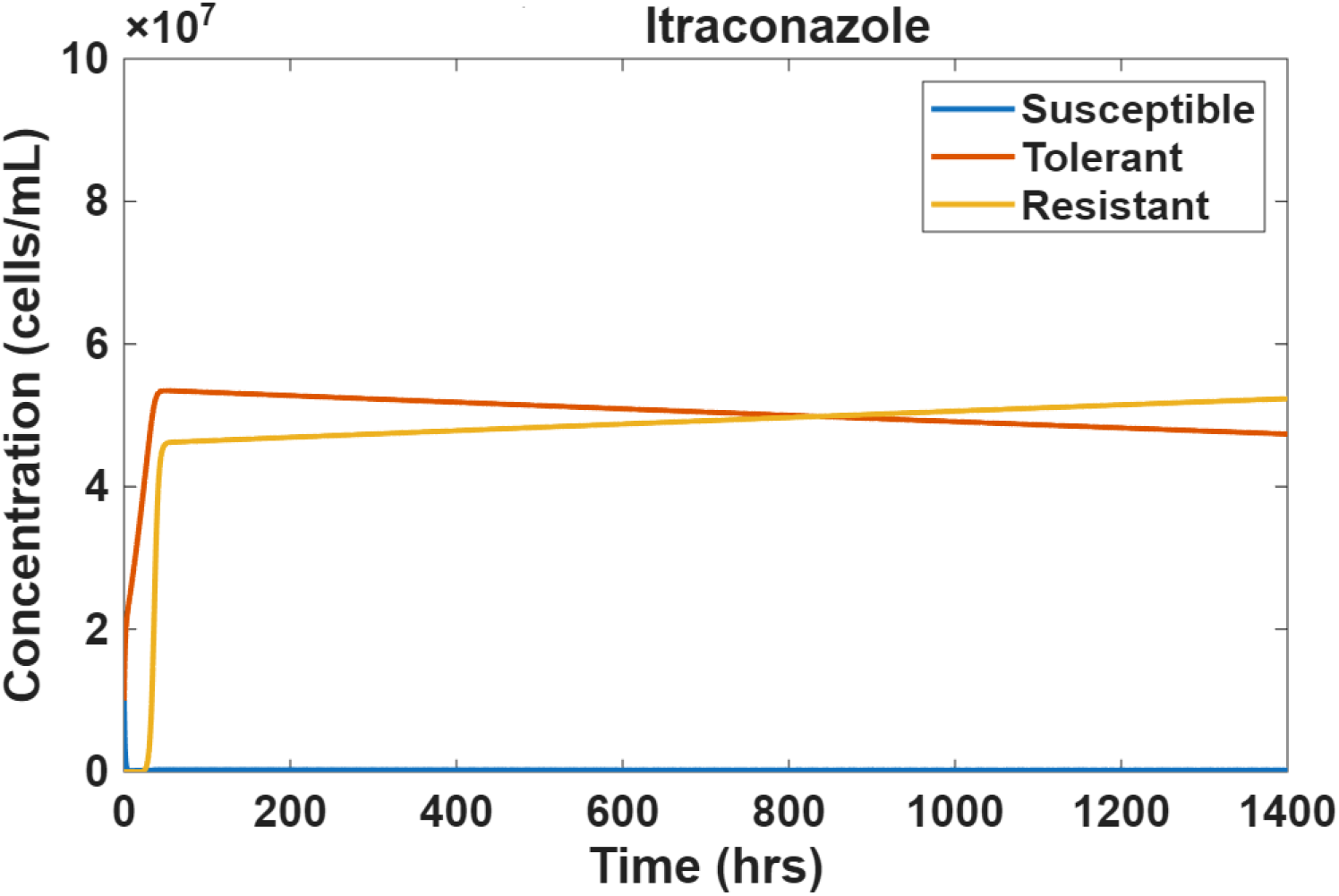
Competition between susceptible, tolerant, and resistant *C. auris* subpopulations during high initial infection and minimum innate immunity. The tolerant subpopulation dominates the population during treatment with itraconazole (3 *µ*g/mL). At approximately 850 hrs, the resistant subpopulation becomes dominant thereafter. High initial infection load corresponds to 10^7^ cells/mL and minimum innate immunity corresponds to 10^4^ cells.

